# Intraspecific variation in the placement of campaniform sensilla on the wings of the hawkmoth *Manduca sexta*

**DOI:** 10.1101/2023.06.26.546554

**Authors:** Kathryn E. Stanchak, Tanvi Deora, Alison I. Weber, Michelle K. Hickner, Abna Moalin, Laila Abdalla, Thomas L. Daniel, Bingni W. Brunton

**Affiliations:** University of Washington, Department of Biology, Seattle, WA; University of Washington, Department of Mechanical Engineering, Seattle, WA

## Abstract

Flight control requires active sensory feedback, and insects have many sensors that help them estimate their current locomotor state, including campaniform sensilla, which are mechanoreceptors that sense strain resulting from deformation of the cuticle. Campaniform sensilla on the wing detect bending and torsional forces encountered during flight, providing input to the flight feedback control system. During flight, wings experience complex spatio-temporal strain patterns. Because campaniform sensilla detect only local strain, their placement on the wing is presumably critical for determining the overall representation of wing deformation; however, how these sensilla are distributed across wings is largely unknown. Here, we test the hypothesis that campaniform sensilla are found in stereotyped locations across individuals of *Manduca sexta*, a hawkmoth. We found that although campaniform sensilla are consistently found on the same veins or in the same regions of the wings, their total number and distribution can vary extensively. This suggests that there is some robustness to variation in sensory feedback in the insect flight control system. The regions where campaniform sensilla are consistently found provide clues to their functional roles, although some patterns might be reflective of developmental processes. Collectively, our results on intraspecific variation in campaniform sensilla placement on insect wings will help reshape our thinking on the utility of mechanosensory feedback for insect flight control and guide further experimental and comparative studies.

## Introduction

Controlled, active aerial locomotion requires sensory feedback to monitor and regulate motor output. For both animals and engineered systems, both vision and mechanosensing play crucial roles in flight control. Mechanoreception is particularly important given its low latency and general utility for rapidly sensing gravity, wind, and structural deformations. Interestingly, the wings of flying animals are not only effective propulsors; they also serve a sensory function, and sensory information from wings feeds back into the flight control system (e.g., Pringle, 1957; Palka et al., 1986; Ando et al., 2011; Fabian et al., 2022).

One type of sensor expected to be important for feedback control during flight in insects is the campaniform sensillum (plural campaniform sensilla, CS), which is a mechanoreceptor that senses deformation in the cuticle resulting from mechanical stress produced by aerodynamic and inertial forces. CS are also found on other parts of the insect body (Snodgrass, 1935), including the legs (Pringle, 1938b), mouthparts (Pringle, 1938a), and antennae (Gewecke, 1972). A CS is a dome-shaped sensor, where the dendrite of a single bipolar neuron innervates a cuticular dome suspended over a cuticular depression (Snodgrass, 1926). Local deformation of the wing results from the interaction between the structural mechanics of the wing with the inertial and aerodynamic forces acting on the wing, causing a local strain field. This force results in movement of the CS dome relative to the surrounding cuticle, causing the neuron to fire. Moreover, the observation that CS neurons on the wing project to the ventral nerve cord in the pro-, meso- and metathoracic ganglia as well as to the subesophageal zone of the brain (Nardi, 1983; Ando et al., 2011) suggests they play a role in proprioceptive feedback.

Depending on the insect species, wing CS can be distributed over the entire wing (but almost exclusively on wing veins), on both the dorsal and ventral sides of the fore- and hindwing (reviewed by Aiello et al., 2020). Although each campaniform sensillum can only sense local strain at the location of the sensor, a distribution of CS provides a representation of the strain distribution across the wing surface. Specifically, rotations of the body about different axes cause characteristic patterns of wing bending and torsion (Eberle et al., 2015). Accurately sensing the resultant strain patterns could allow the insect to estimate and then compensate for any undesired rotation (Mohren et al., 2018). Further, experimentally induced wing bending produces reflexive abdominal flexion in moths, suggesting that wing strain sensors indeed modulate flight reflexes (Dickerson et al., 2014).

Each CS is believed to respond to a characteristic phase of the wingbeat, and thus a wing sensillum functions as a strain-event sensor during flight (Dickinson, 1990; Yarger and Fox, 2018; Gettrup, 1965). Because each CS responds to local strain, its location on the wing impacts its phase selectivity and its sensitivity to perturbation. In flying insects, dense fields of small CS are found at the wing base, where strain is high (e.g., Aiello et al., 2020; Agrawal et al., 2017; Gnatzy et al., 1987; Dickinson, 1990; Cole and Palka, 1982). However, in many insects, individual CS are also sparsely dispersed across the wing veins, distal to the base. The pattern of this sparse distribution of CS varies across the few flying insect species for which wing CS locations have been mapped (Aiello et al., 2021b).

Even more fundamentally, it is unclear if, or how, the distribution of CS varies among individuals of the same species, because most mappings of wings have considered only one specimen of a species. Based on mappings from different authors in limited species, there is some indication that placement can indeed vary among individuals (e.g., slight differences in the mapping of (Dombrowski UJ, 1991) and (Dickerson et al., 2014) of the wing of the tobacco hawkmoth *Manduca sexta*). Understanding variation in CS placement may provide clues to where strain sensing is most critical, in the context of morphological and behavioral characteristics of a particular species. Variation in placement may also indicate how robust strain sensory feedback integration is to differences in available sensory information.

Our goal in this study was to measure intraspecific variation in campaniform sensilla (CS) placement on the wings of *Manduca sexta*. We focused on the broad, distal planform areas of both the dorsal and ventral surfaces of both the fore- and hind-wings, where CS are generally found individually or in pairs. We found that in *M. sexta*, the exact placement of CS varies considerably among individuals. While CS are generally consistently found on the same veins across individuals, there is often wide variation in how those CS are distributed along a specific vein. Thus, here we describe previously unrecognized variation in CS placement across insect wings and discuss the implications of this variation for understanding strain sensory integration in feedback control of insect flight.

## Methods

### Specimen preparation

We collected 9 forewings and 10 hindwings from a laboratory colony of *M. sexta* based at the University of Washington. All forewings and hindwings were from different individuals, but we did not take both the forewing and the hindwing from the same individual. We collected either the right or left wing. As a consequence, we analyzed the forewings and hindwings separately and did not look for any relationship between forewing CS placement and hindwing CS placement. We clipped each wing slightly distal of the base so that the wing could lie flat. We gently manually descaled each wing using soft wipes and forced air, then mounted the wing between two large glass coverslips held together at the edges with glue or tape.

### Locating Sensors

We used the roughly circular cuticular dome as a distinguishing feature of CS to identify them under brightfield light microscopy. In addition, the cuticle of the dome fluoresces under ultraviolet excitation (Thurm, 1964; Michels and Gorb, 2012) (Fig. 1a & b). We took advantage of both of these features to localize CS on the wing surfaces using the 10x objective on a Nikon Ti-2E microscope in both brightfield and with a low-wavelength filter (excitation 340-380, dichroic 400, emission 435-485; i.e., the “DAPI” range). In addition, we also found several CS oriented on the sides of veins, such that the sensillum dome points toward the leading or trailing edge of the wing (Fig. 1c & d).

**Figure 1:**
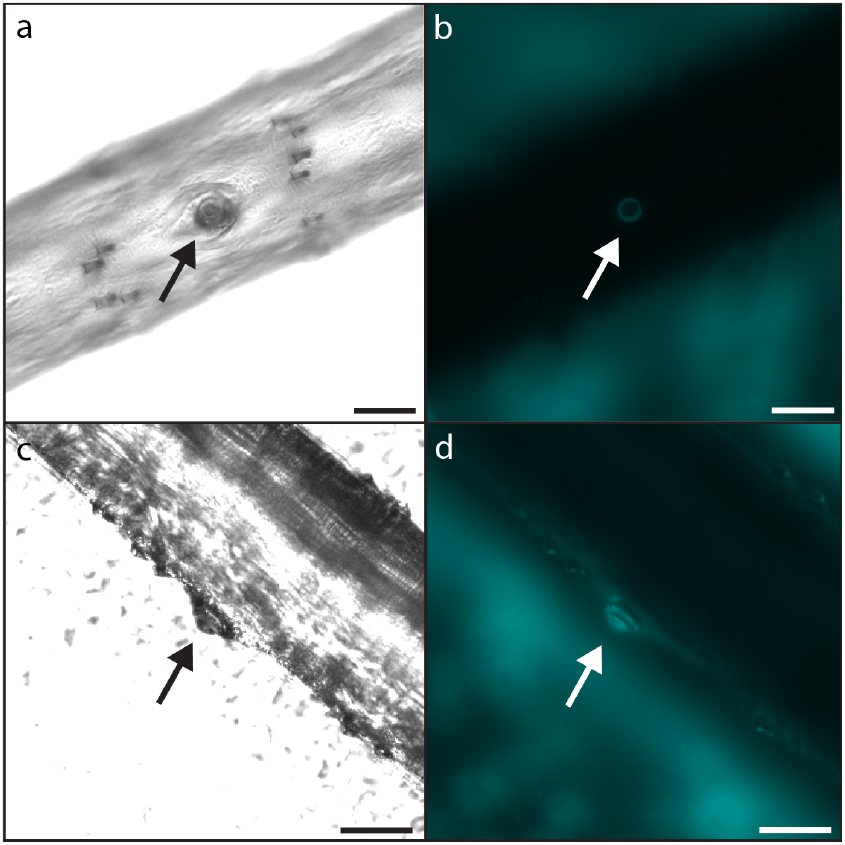
Campaniform sensilla (CS) on moth wings are visible in brightfield (a) and are even more apparent under low-wavelength illumination (b), which makes the cap and collar readily visible due to autofluorescence of the wing cuticle. CS can be found on the top of veins (a and b) or on the sides of veins (c and d), such that they are oriented to face either the leading or trailing edge of the wing rather than face orthogonally to the plane of the wing (as in a and b). Scale bars are 50 *μ*m.

Initially, two researchers (AM and KES or LA and KES) independently examined each wing for CS. We generally restricted the search to the wing veins, including the vein edges, although we did spot-check intra-veinal membranes. We found CS only on veins, with a single exception (see Results, Central/Proximal Veins: Dorsal). After we collected preliminary sensor maps, we then did a targeted search of specific wings areas in which no CS were found in one individual but where sensors were found on others. To ensure consistency and minimize systematic error due to human biases, one researcher (KES) examined all wings following the initial mapping. At the proximal end of the wing—where the wing was detached from the wing hinge—the thickness of the veins precluded light transmission and thus limited our ability to consistently detect sensors. We performed all microscopy using the Nikon software NIS-Elements.

To accurately record the location of each sensillum in relation to the full wing, we first centered each sensillum under the 10x objective and imaged it in both the brightfield and fluorescent light channels. These images served as confirmation of a campaniform sensillum and also allowed us to see large-scale difference in CS morphology. Then, without disturbing the position of the wing mount in the stage, we captured a tiled, full image of the wing surface (or of a relevant vein in the case of re-checking) under the 4x objective. Using a recorded X-Y coordinate error between the 4x and 10x lenses, we then mapped the locations of the CS obtained using the 10x objective into the coordinates of the wing image taken with the 4x objective for downstream analyses, using a custom Python script. Diagrams of CS placement on each individual wing are provided in the Supporting Information.

### Normalizing Sensor Locations

All data analysis was completed in R (v.4.2.2) and ImageJ (v.2.3.0) in FIJI. To compare CS locations along the distal veins, we measured the distances from the most distal intersection of a vein to each sensor as well as to the distal tip of the vein. This was done with the CS locations plotted on the image taken with the 4x objective using the *measure* functions and the *segmented line* tool in ImageJ. The total length of the vein (from the most distal intersection of the vein to the most distal tip) was then normalized to unit length, and the distances of each CS along that vein was normalized accordingly. This normalization allowed us to directly compare variation in CS placement along the vein among different individual moths. Our method assumes allometric wing shape variation, which may not be justifiable across wings of very different sizes (like those of different species), but adult *M. sexta* wings are all similar in size. We did not perform this calculation for veins that had a proximal end cut where the wing was removed from the wing hinge (e.g., the anal veins) to avoid distortion due to inconsistent wing removal.

We next analyzed the often-paired CS found at the distal tip of veins on the ventral surface of both the fore- and hindwings. When paired, both CS were close enough to be captured in one 10x microscope image, and we could categorize their placement relative to each other.

## Results

### Distal Veins: Dorsal

Campaniform sensilla (CS) are consistently found distally on the same longitudinal veins (medial 1, M1; medial 3, M3; cubital 1, CU1; and the anal veins) across individuals (according to the Comstock-Needham system of vein nomenclature; Comstock (1918); Fig. 2 and 3). This is true for both the forewings and the hindwings; that is, the veins on the hindwing that have CS are the same as those on the forewing that have CS. That is, the putatively serially homologous veins of the forewing and hindwing are the veins with CS. And, the same distal longitudinal veins have both dorsal and ventral CS.

**Figure 2:**
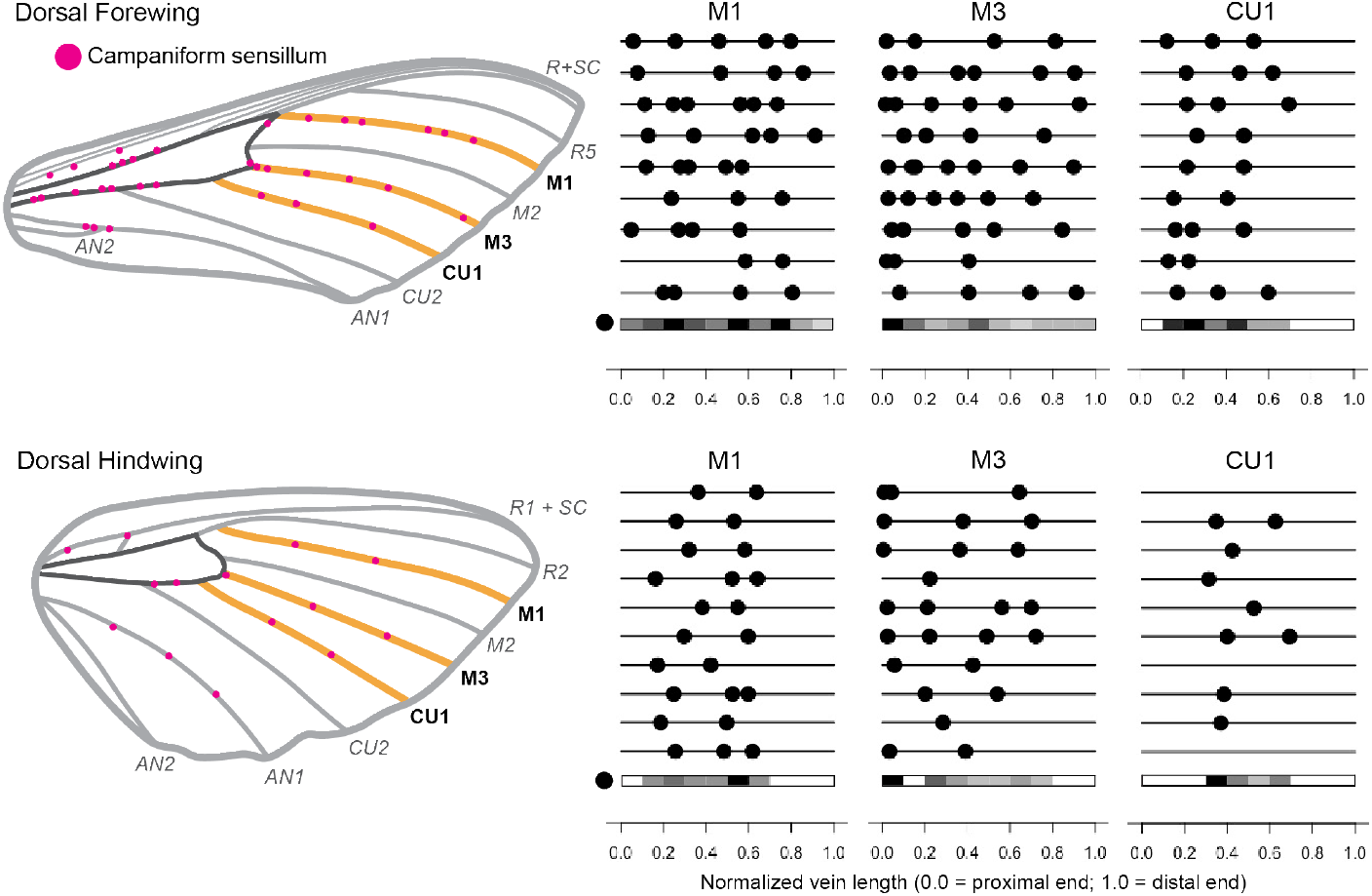
Campaniform sensilla (CS) are generally found on the same dorsal veins of *Manduca sexta* wings across individuals, although the distribution of sensilla along those veins varies. Wing schematics show locations of dorsal CS on one example individual. The veins around the central cell are dark grey, while the veins that correspond to the line diagrams are in gold. The line diagrams next to each wing image demonstrate the distribution of CS on M1, M2, and CU1 for different individuals (9 forewings, 10 hindwings). The gray-scale heatmaps below each line diagram for each vein demonstrate where CS are commonly (or not commonly) found along the length of the vein, split into 10 bins. Black indicates the location with the most CS (for that particular vein), while white indicates no CS.

**Figure 3:**
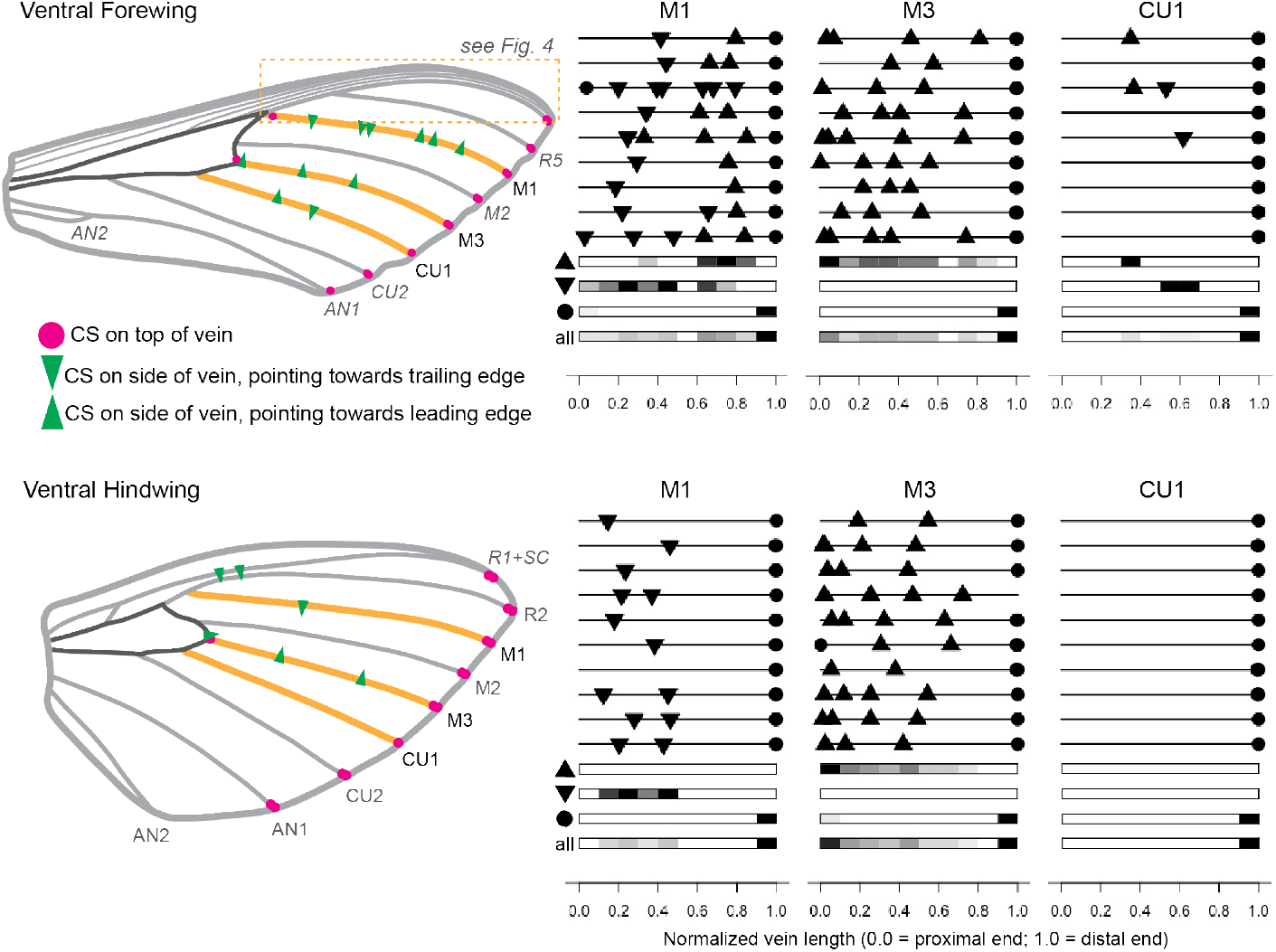
Like the dorsal campaniform sensilla (CS), ventral CS are generally found on the same veins of *Manduca sexta* wings across individuals, although their distribution varies. Wing schematics show locations of CS on one example individual. The veins around the central cell are dark grey, while the veins that correspond to the line diagrams are in gold. The line diagrams next to each wing image demonstrate the distribution of CS on M1, M3, and CU1 for different moth individuals (9 forewings, 10 hindwings). The ventral surface of the wing has many campaniform sensilla that are located on the sides of veins (indicated by green triangles), such that their dome points roughly toward the leading or trailing edge of the wing (direction indicated by triangles). Below each set of distribution plots are heatmaps of CS locations along that particular vein, where the length of vein is split into 10 bins. Black indicates the location with the most CS (for that particular vein), while white indicates no CS..

Although CS are consistently found on the same distal veins, the distribution of CS across these veins is not consistent across individuals (Figs. 2 & 3). The total number of CS on each vein and their placement along the vein both vary. Despite this variability, there are nevertheless some general trends. On the dorsal forewing, CS are distributed across M1 and M3 but are concentrated somewhat more densely on the proximal end of M3. CS are also concentrated more heavily towards the proximal side of CU1. On the dorsal hindwing, CS are distributed in the proximo-medial part of M1 and M3, although several wings have CS very proximal on M3. On CU1, most individuals have only one or two CS located in the middle and none in the far proximal or far distal regions. Not all hindwings have CS on the dorsal side of CU1 (3/10 do not).

### Distal Veins: Ventral

Along the ventral side of the distal longitudinal veins, we found several sensors oriented on the sides of the veins, such that the full cap of the sensillum was not visible under the microscope (Fig. 1c & d). We were still able to distinguish the cuticular cap and collar of CS under low-wavelength light and positively identify them as CS by comparing with other structures visible on the wing. We did not find these on the along the dorsal side of the distal longitudinal veins. The side of the vein on which these CS are found is consistent based on their placement; for example, CS are on the side of M3 facing the leading edge of the wing. On the forewing M1, proximal CS face the trailing edge of the forewing, while distal CS face the leading edge. CS on the sides of the hindwing M1 face the trailing edge of the hindwing. Only a few individuals had CS on the sides of CU1. These were always on the forewing, and this small sample followed the opposite pattern of those on M1; the proximal CS faced the leading edge with those on the distal half of the vein faced the trailing edge. In the forewing of *M. sexta*, the first radial (R) branches, the subcosta (SC), and the costa (C), are very close to one another, forming a very stiff leading edge of the wing. We found numerous CS along these veins on the ventral side of each forewing we examined (Fig. 4). Due to the closeness of the veins, the varied orientation of the sensors, and the high number of sensors, we found it difficult to ensure we located every sensor. Here we only report the presence of these sensors and note that there is some variation among individuals, but we did not quantify their distribution. To our knowledge this is the first report of CS on the ventral leading edge of the forewing of *M. sexta*.

**Figure 4:**
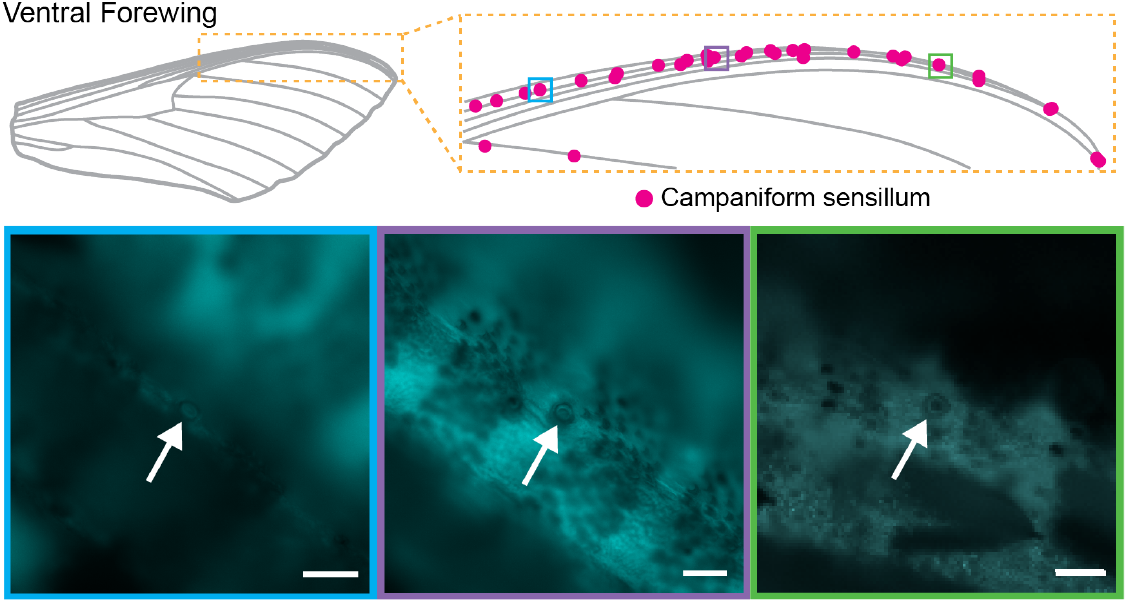
There are many campaniform sensilla (CS) distributed across the leading edge of the ventral forewing. This is a representative distribution from one wing, with magnified views of three CS within this distribution taken under low-wavelength light. Scale bars are 50 *μ*m. In the distribution diagram, each CS is noted with a single dot regardless of where it is located on the vein (top or side). In the microscope images, the CS are indicated with arrows.

The longitudinal veins on the ventral sides of the wing always have two CS (most often) or a single CS at the very distal tip (Fig. 3 and Fig. 5a & b). Whether one or two CS are present depends on the vein and to some extent, the individual. On the forewing, we only found a single CS at the distal tip on AN2, CU1, and CU2—the veins closest to the trailing edge of the wing—in some individuals (Fig. 5c). Even so, most individuals had paired CS at the ends of these forewing veins. Single CS at the distal tips of ventral veins are more frequently found on the hindwing, especially on M3 and CU1.

**Figure 5:**
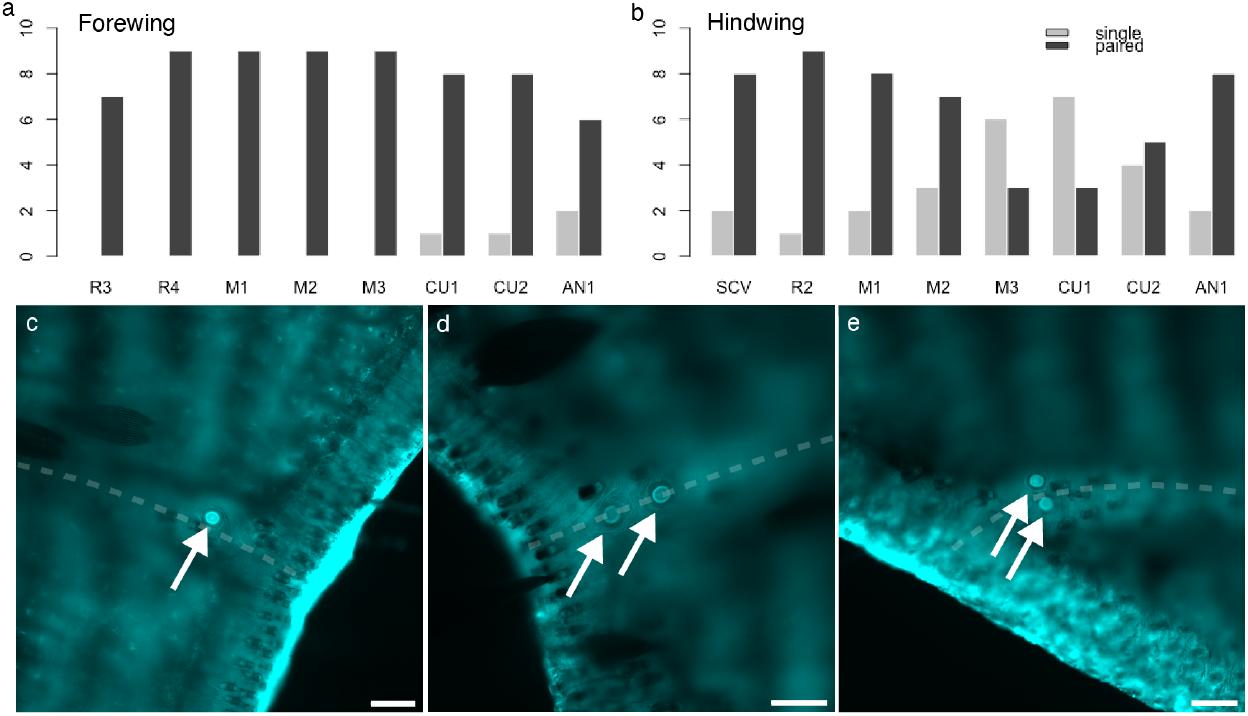
Campaniform sensilla at the distal tip of the ventral longitudinal veins are most often found in pairs, but they also occur singly. Barplots show the number of individual wings with a single or a paired CS on the tips of the longitudinal veins, ordered from leading-to-trailing edge of the wing, for the forewing (a) and the hindwing (b). Some sample wings were damaged at one or more vein tip such that we could not confirm the number of CS; thus not all bars sum to the total sample of wings. Images of CS are of a single sensillum at a vein tip (c), a paired set aligned with the axis of the vein (d), and a paired set aligned off-axis of the vein (e). The dashed line in (c-e) roughly follows the longitudinal axis of the vein. Scale bars are 50 *μ*m.

**Figure 6:**
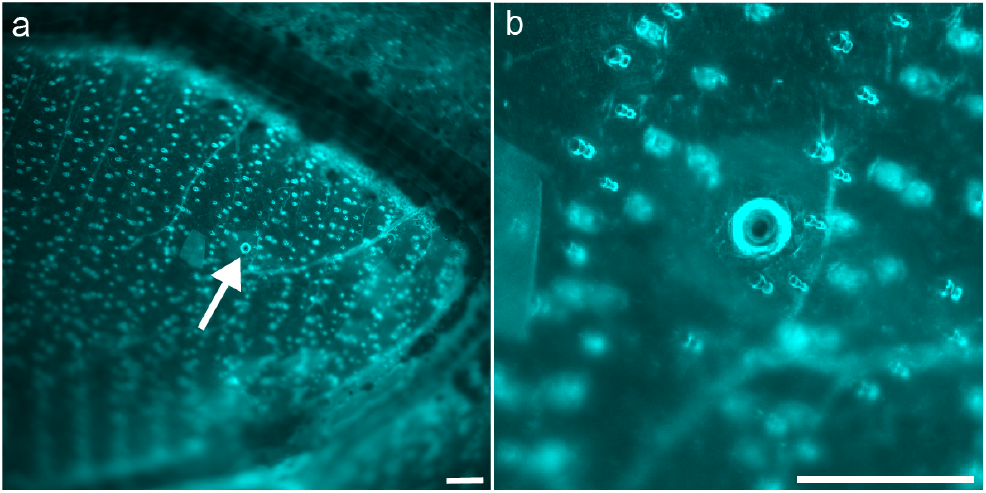
We found one feature resembling a campaniform sensillum that was located on the wing membrane in-between veins, rather than on a vein itself (a). A closer image of the sensillum-like structure is in (b). Scale bars are 100 *μ*m.

When the CS are present as a pair, they are generally aligned serially, such that the vector between the two sensors is roughly parallel with the longitudinal axis of the vein (Fig. 5d). There are occasional exceptions, most notably on AN2, where the two CS are instead beside each other, such that the axis between the sensors is roughly perpendicular to the long axis of the vein (Fig. 5e).

### Anal veins

The number of CS on the dorsal anal veins varies among individuals. On the dorsal anal veins of the forewing, we found between two and six CS distributed on AN1, near the junction of AN1 and AN2. On the dorsal forewing AN2, we found between zero and three CS, although these were difficult to distinguish from other wing features. On the dorsal anal veins of the hindwing, we found between two and six CS distributed across AN1, except for one individual on which did not find any. We did not find any CS on the dorsal hindwing AN2. On both the forewing and the hindwing, the only CS on the ventral side of the anal veins were the paired or single CS at the very distal end of AN1.

### Central/Proximal Veins: Dorsal

In *M. sexta*, the pre-branching radius and media form a central cell, closed at the distal end by a cross-vein that connects M1, M2, and M3 (Fig.2 & 3). The proximal ends of veins forming this cell were generally too thick to permit enough light transmission for accurate CS identification, but we could positively identify CS more distally.

On every dorsal forewing, we found several CS on both sides of the base of the central cell (i.e., at the media and radial bases). On at least five wings, we also found many CS at the base of the SC. These proximal ends were quite dark and additionally had more wing features like scale sockets, and we could not verify the presence or number of CS at these locations on every wing. Seven of the dorsal forewings had two CS on the media between the intersections with CU1 and CU2, while the remaining two forewings had one but a second at the base of CU2. Six of the dorsal forewings had one CS at the base of M3 on the cross-vein. As noted above, M3 also commonly has at least one CS at the proximal end of M3 (Fig. 3). On two dorsal forewings we found a CS near the base of M1, on the cross-vein between M1 and M2.

Similar to the forewing, on every dorsal hindwing we found at least one CS near—but just distal to—the base of CU2, and on six we found another CS along the media between CU2 and CU1. On several wings we could distinguish CS proximal of CU2, but the wing was often too thick to make confident determinations. The distal longitudinal vein M3 often had at least one CS at its proximal end near where it meets the cross-vein of the central cell (see Fig. 2), but in one specimen we found one CS near the base of M3 but on the veins around the central cell between M3 and CU1. On two dorsal hindwings we found a CS on the cross-vein between the base of the R2/M1 branch and M2. Interestingly, on one individual wing we also identified a campaniform-like structure on the interveinal wing membrane in about this same location—near the base of the R2/M1 branch and M2 (Fig.6).

### Central/Proximal Veins: Ventral

There were few CS around the proximal veins surrounding the central cell on the ventral forewing. On seven ventral forewings, there was one CS on the cross-vein near the base of M3.

On eight ventral hindwings, there were two CS near the midline of the SCV oriented such that they faced the trailing edge of the hindwing (see example in Fig. 3). On one other hindwing there was only one CS in this location, while on the last we did not see any, although they could have been obscured by remnant scales. On seven ventral hindwings, we found one CS at the base of CU2, and on five we found one CS at the cross-vein near the base of M3.

#### Box 1

Trends in placement of campaniform sensilla (CS) on wings of *Manduca sexta*

- On the dorsal wing, CS are found on the same distal veins on both the forewing and the hindwing, but the number and placement of CS along those veins varies among individuals.
- On the ventral wing, CS are often found on the sides of veins. The CS always face the leading edge of the wing on M3, while on M1 they face either the leading edge (distal half of forewing M1) or trailing edge (hindwing M1 and proximal half of forewing M1).
- Several CS are found along on the ventral side of the forewing near the leading edge.
- CS are generally found in pairs at the distal ends of the ventral veins, although on certain veins they can be found singly, as well.
- There is often at least one CS at the proximal base of M3 on the ventral side of both the fore- and hindwings.
- On the hindwing, the ventral side of the vein SC often has a pair of CS that face the trailing-edge of the wing.
- On both the dorsal fore- and hindwing, there are one or two CS on the media outlining the central cell between CU2 and CU1.
- There are more CS around the central cell of the dorsal forewing than the hindwing, and there are fewer on the ventral side of the central cell than on the dorsal side.

## Discussion

In this paper, we found that although campaniform sensilla are found in roughly the same general locations on the wings of *M. sexta*, the number and specific distribution of these sensors are highly variable, so that the idea of homologous individual sensors is largely unfounded in understanding campaniform sensilla placement. Our findings are summarized as trends in Box 1. Our results complement those of a recent study that found interindividual variability in CS placement on the legs and slight variability on the wings of *Drosophila melanogaster* (Dinges et al., 2021). Previously, CS were thought to be found in generally stereotyped locations on *D. melanogaster* and the Drosophilidae (Palka et al., 1986; Dickinson and Palka, 1987). Therefore, when hypothesizing implications of CS for feedback control in flight, we consider the regions of the wing where sensors are consistently found, and not necessarily the precise locations and number of sensors, as clues to the important sensory functions of wing CS.

### Functional Consequences of CS placement

Based on our findings of intraspecific variation in CS placement on several regions on the wings—such as the distal longitudinal veins (Figs. 2 & 3)—the sensory feedback control system must be robust to variation in CS placement on a local scale. This is consistent with findings from wing computational models indicating that sensing accuracy is robust to variation in strain sensor placement in wings with stiffness gradients (Weber et al., 2023). Thus the spatial variation in stiffness found in insect wings (Combes and Daniel, 2003) could endow them with inherent robustness to variation in sensor placement. Inter-individual variability may, however, require flexibility in downstream circuits that integrate signals from these CS in order to compensate for differing CS locations.

Nevertheless, some aspects of CS placement are notably consistent across individuals, which may be indicative of the functional roles of CS signals in sensory feedback control of wings. Even though the number and distribution of CS on the distal longitudinal veins varies, the orientations of the CS across individuals are fairly consistent within a vein. On the dorsal side of both the fore- and hindwings, all sensors point away from the wing surface; that is, the CS are oriented on the top face of their respective veins. On the ventral sides of the wings, CS are oriented on the sides of the veins such that they face either the leading edge (hindwing and forewing M3; distal half of forewing M1) or the trailing edge (hindwing M1 and proximal half of forewing M1).

Where the CS are placed around the vein (i.e., on the top or on the sides) determines which direction of bending or torsion the sensor will be most sensitive to. A flexible insect wing can bend about multiple axes as well as twist (experience both bending and torsion). Fig. 7 demonstrates the pattern of strain (principal strain) that arises at the surface of cylindrical beam in bending. The maximum strain is experienced at the very top and the very bottom of the beam, with a gradient toward the middle, where there is no strain (the neutral axis). Thus, CS located on the top of a vein would respond to different strain directions (and thus different stress application directions) than CS located on the sides of the wing. For example, a CS positioned directly on top of the ventral surface of a vein is likely to be most sensitive to bending of the wing in either the dorsal or ventral direction, as this will cause it to experience the greatest strain (Fig. 7). A CS positioned facing the leading edge on the same vein will instead experience greatest strain when the wing bends towards the leading or trailing edge. CS orientation on the veins may also impact the phase of the wingbeat at which a given CS responds, as CS positioned directly at the top or bottom of a vein would reach its strain threshold at a different phase of the wingbeat than those positioned to the sides.

**Figure 7:**
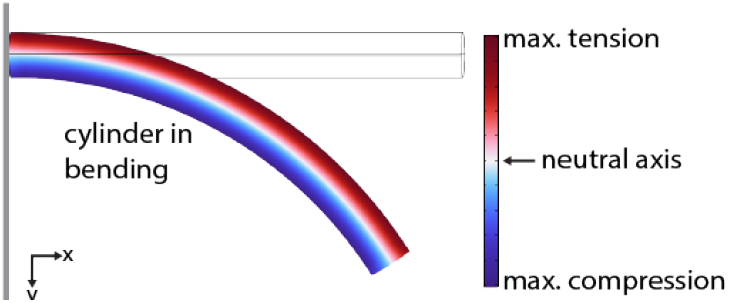
An insect wing vein under loading can be thought of as a cylinder under bending. The color scale represents the x-x normal strain due to the stresses caused by bending. The top of the cylinder experiences the most strain due to tension, while the bottom of the cylinder experiences the most strain due to compression. The transition from tension to compression occurs at what is known as the “neutral axis”, where the strain is zero. The wireframe indicates the at-rest position of the cantilevered cylinder.

The microscopy methods we employed did not allow us to measure the shape of the CS, although we were able to observe that the CS on the tops of the veins were generally circular. Circular CS are thought to respond equally to strain in all directions unlike the elliptical CS that are often found in dense fields at locations like the wing base (Pringle, 1948, 1938b; Zill and Moran, 1981; Dickinson, 1992). In the situation depicted in Fig. 7, assuming the planform area of the wing is facing up out of the page, CS on the leading edge side and the trailing edge vein of the approximately cylindrical vein would experience strain of similar magnitude (tension or compression, respectively). If they are also circular, benefits to being on one side of the wing rather than another may be due to subtle effects related to variable wing stiffness and wing shape asymmetry. If they are elliptical, however, they may be more receptive to tension or compression, depending of the orientation of their major axis. A detailed understanding of the forces experienced by a CS during flight is lacking, and our findings here illustrate the need for computational models of CS in wings, where wing dynamics and the morphology of the sensillum and wing can be manipulated to query how a sensillum responds to local strain. Small changes in patterns of wing thickness or transitions between cells with different material properties can lead to areas of locally increased strain called strain concentrations. Determining the location on the veins subject to strain concentrations requires high-fidelity anatomically accurate measurements or models, such as have been done for a *Sympetrum striolatum* dragonfly hindwing Fabian et al. (2022).

The consistent presence of numerous CS distributed along the leading edge of the ventral forewing was previously unreported in *M. sexta*. The close proximity of these veins results in a stiff leading edge, and the relatively dense distribution of CS along these veins might be critical for detecting slight spatial variations in strain during flight. Vortices produced by a stiff leading edge are thought to play an important role in generating lift (Ellington et al., 1996) (and are modulated by tip vortices in smaller insects; Birch and Dickinson (2001)). Thus wing cuticle strain resulting from leading-edge vortices might be important to detect. Insect wings also twist a lot during flight, resulting in torsion along the leading edge and wing camber (Ennos, 1988; Wootton, 1993), the detection of which could also be critical for wing morphing and flight regulation. In addition, wing stiffness is one determinant of accuracy in sensing rotation (Weber et al., 2021), and nonuniform stiffness further improves the detection of body rotation (Weber et al., 2023).

The paired sensors at the distal ends of the longitudinal veins are reminiscent of the proximal and distal “twinned sensilla of the margin” of *Drosophila*, which have different developmental origins and project to different neural tracts (Blair and Palka, 1985; Palka et al., 1986). They also have different physiological properties: one is tonically active and one is rapidly adapting to a constant strain stimulus (Dickinson and Palka, 1987). The physiological properties of the distal paired CS on *Manduca* wings are unknown, but for many wing configurations, distally located CS are optimal for sensing rotation (Weber et al., 2021), so having multiple response-types in those optimal locations could be beneficial for flight control. Alternatively, these sensors could be physiologically identical and serve together as a way of sensing spatial derivatives. Yet another alternative is that could simply serve to detect collisions, which are experienced frequently by foraging flying insects (Foster and Cartar, 2011).

### Caveats

We should of course be cautious in assigning functional relevance to trends in CS placement. The wing veins—to which CS are typically restricted—have other physiological roles including circulation and respiration (Salcedo and Socha, 2020; Tsai et al., 2020), in addition to providing wing stiffness and innervation, which could constrain CS to non-optimal locations. These multiple constraints could limit the ability of sensory neurons to access all possible sensor placement locations along veins, including potentially limiting which side of the vein the neuron can access. CS placement could also be partially determined by developmental constraints. For instance, that CS are found on the same distal longitudinal veins on the forewing and the hindwing may be a consequence of developmental patterning of the veins and serial homology rather than an indication of functional relevance. In addition, we note that our samples are from a highly inbred colony of moths that has been released from selection pressure for several generations.

Our methods of finding CS via transmitted-light microscopy was adequate for finding CS on the distal, planform surface of the wing, but it was not adequate for confirming CS along thicker veins or where there were more remnant features like remaining scales or large sockets for scales. Because of the large size of the *Manduca* wing, we could confirm and quantify the presence of CS but not their absence. Although we took a considered approach to searching for CS multiple times, we may have missed some CS. Nonetheless, we are confident our method avoided systematic errors (other than where veins were too think for light transmission, as already noted), and thus are appropriate for our reported results. Scanning electron microscopy (SEM; e.g., (Palka et al., 1986; Dickinson and Palka, 1987; Dinges et al., 2021)) is still a better standard for verifying CS morphology and could be used in the future to better characterize the morphology of CS as well as their exact positioning around the circumference of a vein. Another possible method for identifying sensors is through nerve tracing, as done in (Fabian et al., 2022), in which they identified sensors at neuronal terminals and characterized their morphology. The downside of these methods (SEM and nerve tracing) is the time required to prepare specimens to map the sensors. In addition to CS, other mechanosensors are also found on insect wings, including bristle sensors (in which deflection of a hair-like bristle causes a neuron to fire) and several other cell types (see Fabian et al. (2022)).

### Implications for Future Studies

Unlike the reportedly consistent locations of CS on the wing surface of Drosophilidae and other flies (Dickinson and Palka, 1987), CS on the wings of the bombycoid moth *Manduca sexta* display notable variation in number and distribution, even if they are generally found in similar locations. The trends in CS placement noted here (Box 1) could serve as a basis for setting hypotheses and predictions for broader comparative studies, using the Bombycoidea as a model clade, which exhibits variation in size, wing shape, and wing kinematics (Aiello et al., 2020, 2021a). How CS placement varies with morphological, behavioral, and ecological factors could provide clues to the functional role of wing strain-sensing as well as help build an understanding of the many constraints under which the insect wings evolve.

## Supporting information

Supplemental Figure - Hindwings

Supplemental Figure - Forewings

## Acknowledgements

We thank Brett Aiello, Sweta Agrawal, and John Tuthill for discussions about campaniform sensilla placement, and Wai Pang Chan of the UW Biology Imaging Facility who assisted with the microscopy methods. We acknowledge funding support from the Air Force Office of Scientific Research (award FA9550-19-1-0386 to B.W.B. and T.L.D.). L.A. and A.M. were supported by the UW-ENDURE program through the National Institutes of Health Award R25NS114097.

