## Supplementary figures and images for "Intraspecific variation in the placement of campaniform sensilla on the wings of the hawkmoth *Manduca sexta*"

### Supplemental Figure - Forewings

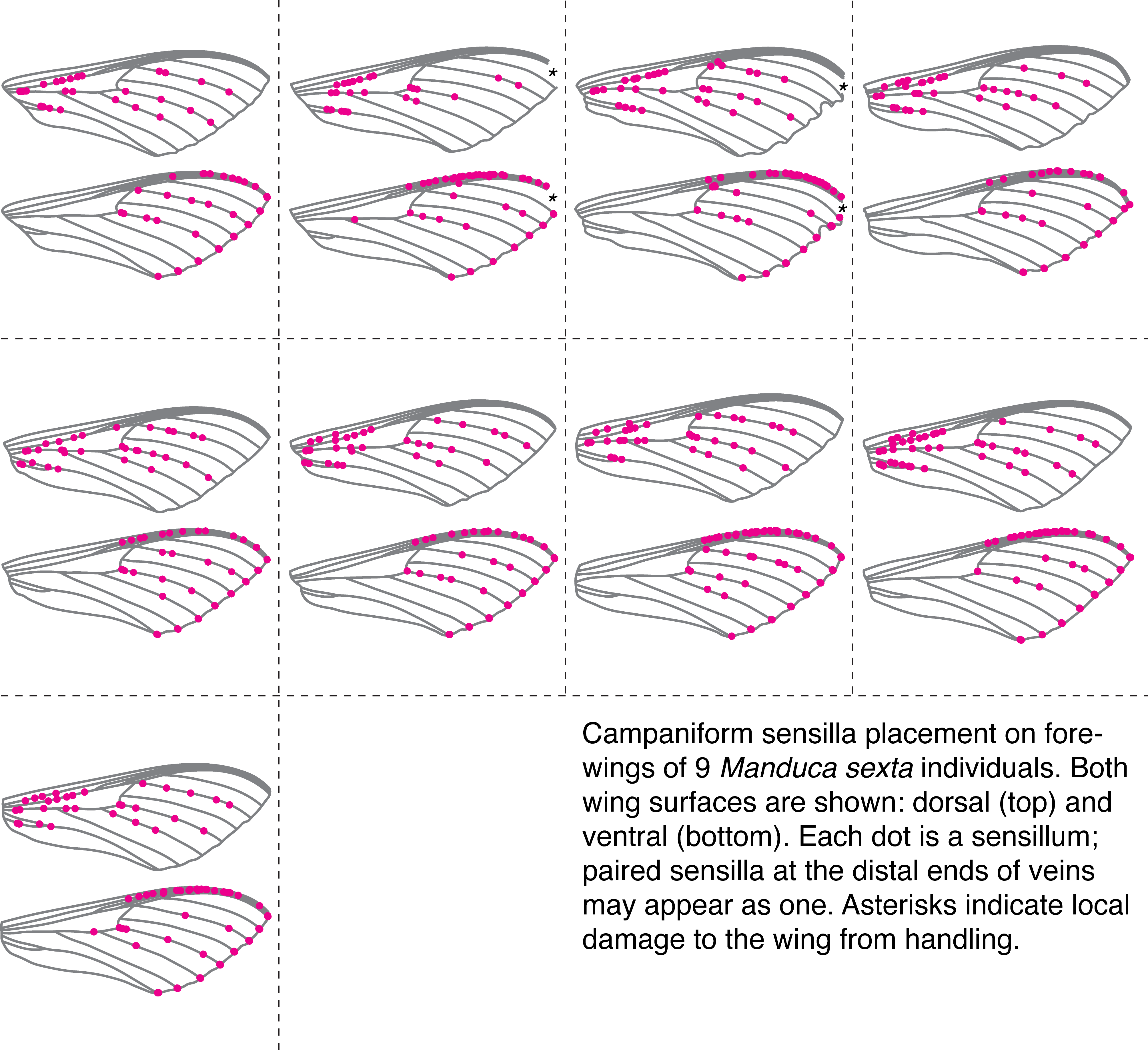

### Supplemental Figure - Hindwings

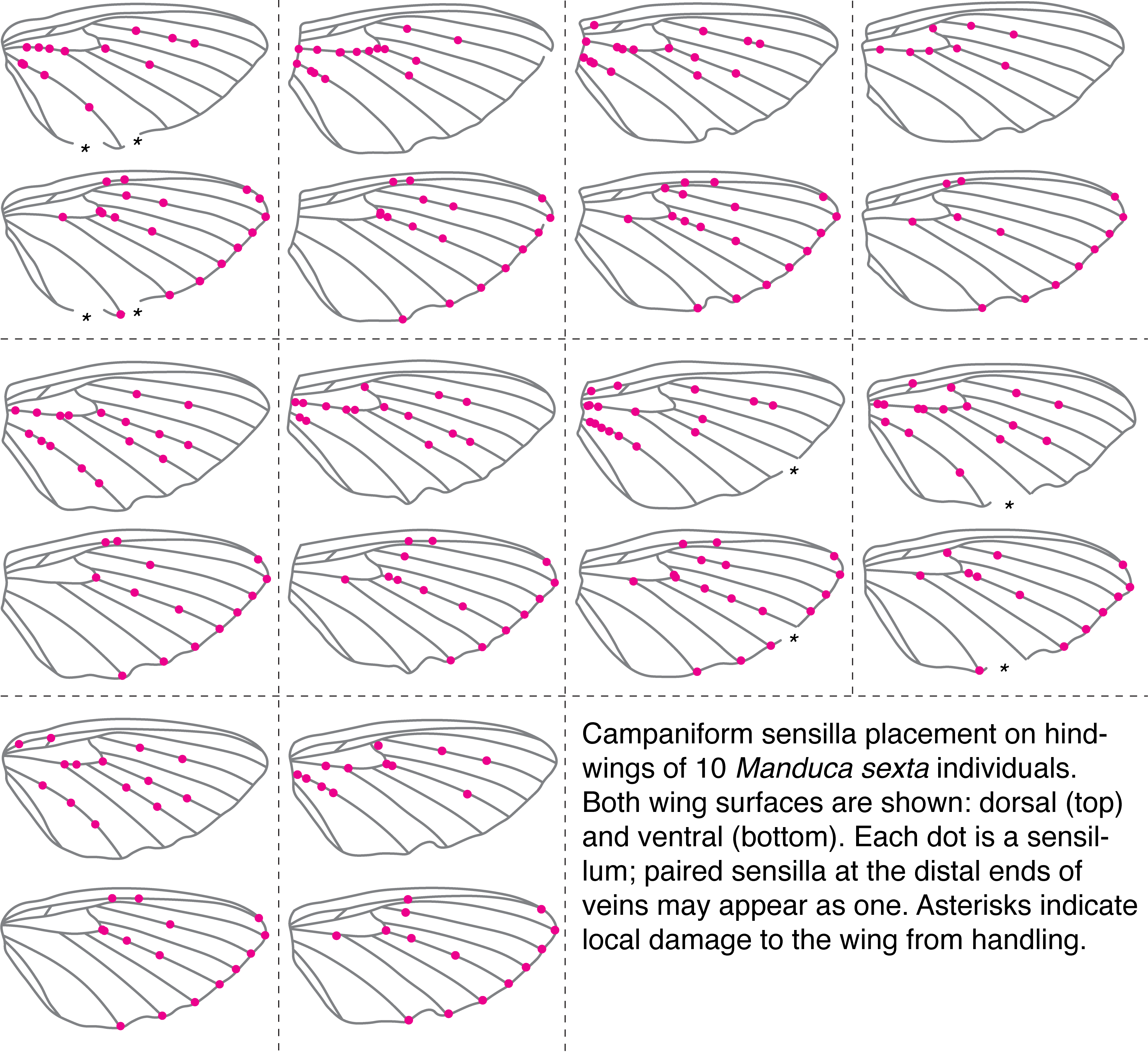
